# Inter-individual Recovery of the Microbiota and Metabolome Over Time Following Fecal Microbiota Transplantation in Patients with Recurrent *Clostridium difficile* Infection

**DOI:** 10.1101/141846

**Authors:** Anna M. Seekatz, Casey M. Theriot, Krishna Rao, Yu-Ming Chang, Alison E. Freeman, John Y. Kao, Vincent B. Young

**Author notes:** **Current addresses:** College of Veterinary Medicine, Department of Population Health and Pathobiology, North Carolina State University, Raleigh, NC. Department of Internal Medicine, Division of Gastroenterology, University of Colorado Anschutz Medical Campus, Aurora, CO. **Address correspondence to Vincent B. Young**.

## Abstract

A significant proportion of individuals develop recurrent *Clostridium difficile* infection (CDI) following initial disease. Fecal microbiota transplantation (FMT), a highly effective treatment method for recurrent CDI, has been demonstrated to induce microbiota recovery, a critical component of disease recovery. However, identification of the specific microbes and their functions that directly impact recovery from CDI remains difficult. We assessed for associations among microbial community members and metabolites in patients with recurrent CDI following treatment with FMT over time to identify groups of bacteria with potential restorative functions. Using 16S rRNA gene-based sequencing, we observed marked similarity of the microbiota between recipients following FMT (n = 6, sampling up to 6 months post-FMT) and their respective donors. Increased levels of the secondary bile acid deoxycholic acid and the short chain fatty acids (SCFAs) butyrate, acetate, and propionate were observed post-FMT. To take into account longitudinal sampling and intra-individual differences, we applied a generalized estimating equation approach to model metabolite concentrations with the presence of specific members of the microbiota. Microbial metabolites that were increased following FMT associated with members classified within the *Lachnospiraceae, Ruminococcaceae*, and unclassified *Clostridiales* families. In contrast, members of these taxa were inversely associated with primary bile acids. The longitudinal aspect of this study allowed us to characterize individualized patterns of recovery, revealing variability between and within patients following FMT.

**IMPORTANCE:** *Clostridium difficile* infection (CDI) is an urgent and serious healthcare-associated problem. In recent years, fecal microbiota transplantation (FMT) has been successfully used to treat recurrent CDI, a frequent outcome of disease. While it is apparent that FMT promotes recovery of the microbiota, it is unclear how microbes and their functions promote recovery from disease. This study aimed to identify associations among microbes and metabolites following FMT and to identify critical microbial functions following FMT treatment for recurrent CDI. Overall, recovery of the metabolome was highly dynamic and individualized in all patients, who were all successfully treated. Our results suggest that microbial changes following FMT may be highly specific to the donor-recipient relationship. Further understanding of the host-microbe environments necessary to enable successful transplantation of microbes during FMT could aid development of specific microbial therapeutics for recurrent CDI and other gastrointestinal diseases.

## Introduction

*Clostridium difficile* infection (CDI) is a significant nosocomial infection and is estimated to cause 453,000 infections a year in the United States (1). Manifestations of CDI can range from mild diarrhea to the more severe pseudomembranous colitis, and recurrence of infection following successful antibiotic treatment or primary disease impacts 20–30% of patients (2). Susceptibility to initial infection is correlated with a loss in diversity of the indigenous gut microbes, the microbiota, such as following antibiotic use (3). Similarly, it is hypothesized that attenuated recovery of the microbiota following antibiotic use contributes to the development of recurrence (4, 5).

The best evidence that a diverse microbiota is required for recovery from CDI is the successful outcome of fecal microbiota transplantation (FMT), with a reported effective rate of 90% in treating recurrent CDI (6). FMT has been used successfully to treat CDI by restoration of microbial populations (old pubs). Given the availability of high throughput sequencing methods to analyze the microbiota in deeper detail, studies comparing the microbiota pre- and post-FMT have demonstrated recovery of overall microbial community diversity and various members of Bacteroidetes and Firmicutes (7–11). However, there is still a lack of identification of specific microbes that directly mediate clearance of *C. difficile* and subsequent recovery from CDI. Furthermore, inherent variability and uniqueness of the gut microbiota in the human population complicates identification of distinct groups of microbes that facilitate recovery from *C. difficile* after FMT, whether occurring by direct colonization from the donor or by indirectly influencing the gut environment.

The changes in microbial communities that lead to susceptibility to CDI are thought in part to be due to changes in metabolites produced by the gut microbiota (12, 13). *C. difficile* exists as a spore in the environment, aiding its transmissibility in hospitals, and disease initiation in its host requires germination. In vitro, it is known that primary bile salts, such as taurocholate, along with glycine are required for *C. difficile* germination (14, 15). In contrast, secondary bile acids, which are products of microbial metabolism, have been shown to inhibit germination, growth, and toxin activity (16, 17). It is known that the intestines of mice treated with antibiotics contain high levels of primary bile salts, associated with susceptibility to CDI (18). More recently, the 7α-dehydroxylase-containing *Clostridium scindens*, which can dehydroxylate the primary bile acid cholate into the secondary bile acid deoxycholic acid (DCA) (19), has been demonstrated to attenuate colonization by *C. difficile* in mice (20). Measurements of secondary bile acids in pre- and post-FMT fecal samples of patients with recurrent *C. difficile* support this observation (21). However, other studies in mice suggest that the provision of dehydroxylation alone does not necessarily mitigate resistance to or clearance of *C. difficile*. Recently, Studer et al demonstrated that mice colonized with a defined community of 14 members, including *C. scindens*, only delayed onset of disease following infection with *C. difficile* (22). Lawley *et al*. have previously demonstrated that a six-member community of bacteria that did not contain a bacterium with 7-alpha dehydroxylase activity was able to clear *C. difficile* in a murine model, resulting in a shift in short chain fatty acids (SCFAs) (23). Furthermore, levels of secondary bile acids in patients treated with FMT, while increased overall, do not necessarily predict failure of FMT to clear *C. difficile* (24). It is likely that the host response as well as additional metabolic pathways, provided by a healthy community with bacteria other than *C. scindens*, also contribute to clearance of *C. difficile* and subsequent recovery from CDI.

This study analyzed the recovery of the microbiota community and metabolic environment in patients with recurrent CDI treated with FMT over time. We aimed to 1) quantify levels of SCFAs in addition to bile acids following FMT and 2) individualized patterns of microbial and metabolite recovery following FMT. We observed increases in SCFAs and secondary bile acids, indicative of restoration of a metabolically active microbiota community.

## Results

### Fecal microbiota transplantation results in diverse, patient-specific microbiota community profiles

Six patients with recurrent CDI successfully treated with FMT, or recipients, provided multiple samples pre- and post-FMT (Figure 1A). Two samples were collected prior to the FMT procedure (pre-FMT: 1–2 weeks and 1–2 days), and 4 samples were collected following the FMT procedure (post-FMT: 1–2 days, 1–2 weeks, 1 month, and 6 months). We obtained the fecal sample used for the procedure from five donors. One aliquot from this donor sample was immediately frozen (unprocessed) and one aliquot was collected following dilution and filtration of the material for FMT (processed) to compare the impact of processing on the sample input. The PCR-amplified V4 region of the 16S rRNA gene was sequenced to investigate the microbiota community in FMT donors and recipients (Table S1).

**Figure 1.**
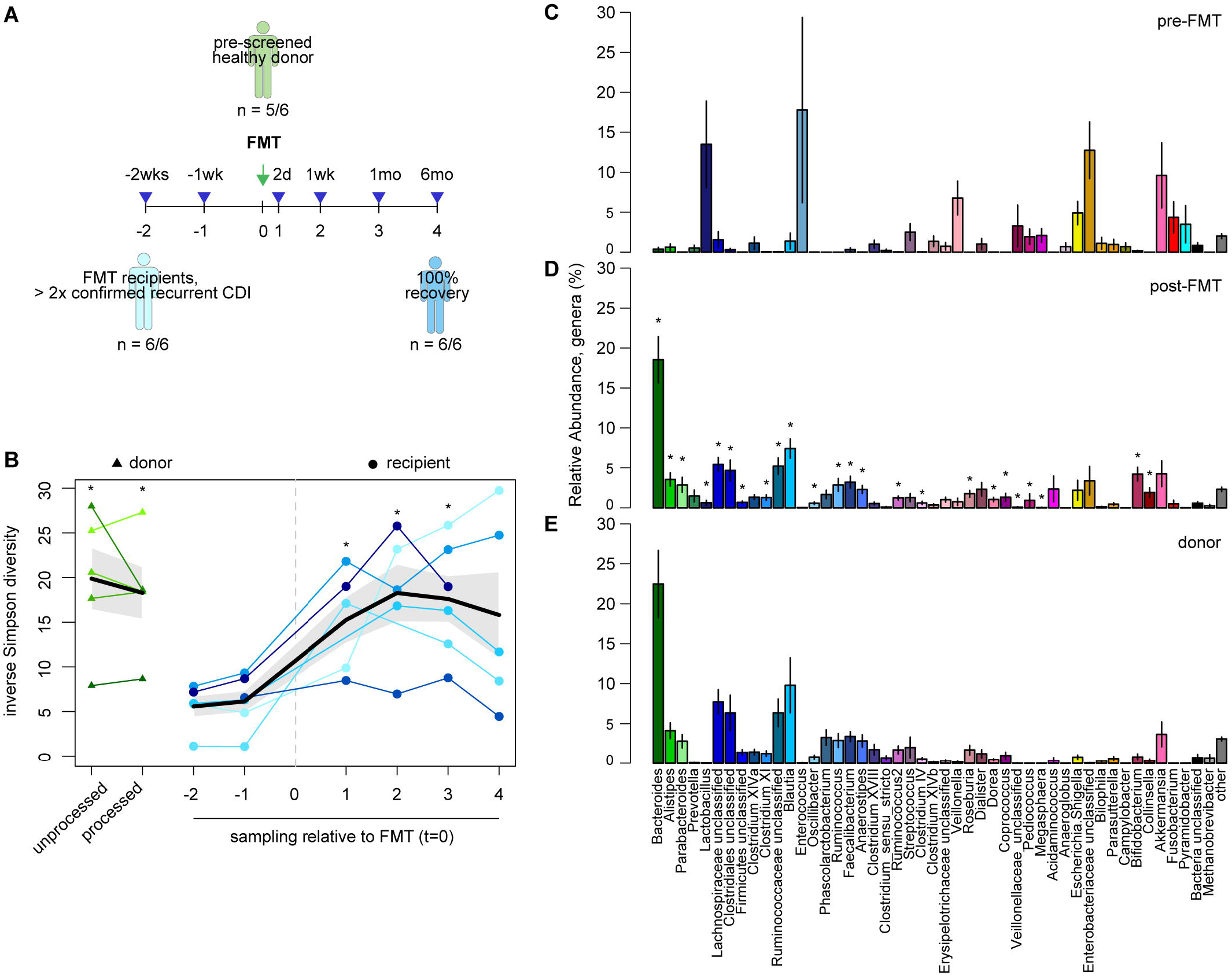
**A**) Study design and sampling scheme. **B**) Fecal microbiota diversity (inverse Simpson index) in FMT donors (unprocessed and processed samples of donated material) and over time in FMT recipients. Individual values are shown per patient (green = donors; blue = recipients), with mean (black line) and standard deviation (grey area) (Wilcoxon test between unprocessed donor sample and recipient time points, p < 0.05). **C-E**) Mean relative abundance and standard error of most abundant genera in recipients pre-FMT time points (**C**) and post-FMT time points (**D**) compared to donors (**E**). Stars (*) indicate significant differences in abundance between pre-FMT and post-FMT time points (Wilcoxon test, p < 0.005).

As expected, the microbiota in donor fecal samples was diverse (inverse Simpson index), with a predominance of bacteria in the Firmicutes and Bacteroidetes (Fig. 1B and E). FMT processing did not impact the measured diversity (Wilcoxon test, p = 0.513). Similar to previously reported findings (10), the recipient patients with recurrent CDI had a fecal microbiota with low diversity prior to FMT (Fig. 1B), encompassing a limited number of bacterial genera, with *Lactobacillus, Enterococcus*, and unclassified Enterobacteriacaeae groups being the most abundant (Fig. 1C, Fig. S1). Following FMT, the microbiota in recipients exhibited a significant increase in diversity compared to their pre-FMT state and remained consistently diverse over time (Wilcoxon test, p < 0.05, Fig. 1B). Taxonomic classification to the genus-level revealed similar community profiles to the respective donor (Fig. 1D, Fig. S1). In particular, levels of *Bacteroides*, unclassified Lachnospiraceae and Clostridiales increased in recipients following FMT (Fig. 1E).

Short sequences of the 16S rRNA gene cannot distinguish species- or strain-level differences between sample comparisons. However, the presence of unique operational taxonomic units (OTUs) across multiple samples within an individual suggests that 16S rRNA gene-based sequencing has the capacity to distinguish members of the microbiota unique to patients. Overall, recipients were more similar to themselves post-FMT than to their pre-FMT state, their respective donors, or random donors and other recipients, post-FMT (Kruskal-Wallis test, p < 0.0005) (Fig. 2A). We compared the number of shared OTUs between recipients and their respective donors to recipients and random donors to distinguish whether specific members of the donor microbiota were colonizing the recipient (patients R1-R5, since no donor sample was available for patient R6; Figure 2B). Recipients were more likely to be similar to their donor than other donors over time, suggesting that at least some microbes may be directly colonizing from the donor, as discriminated by the presence of unique OTUs within donor-recipient pairs.

**Figure 2.**
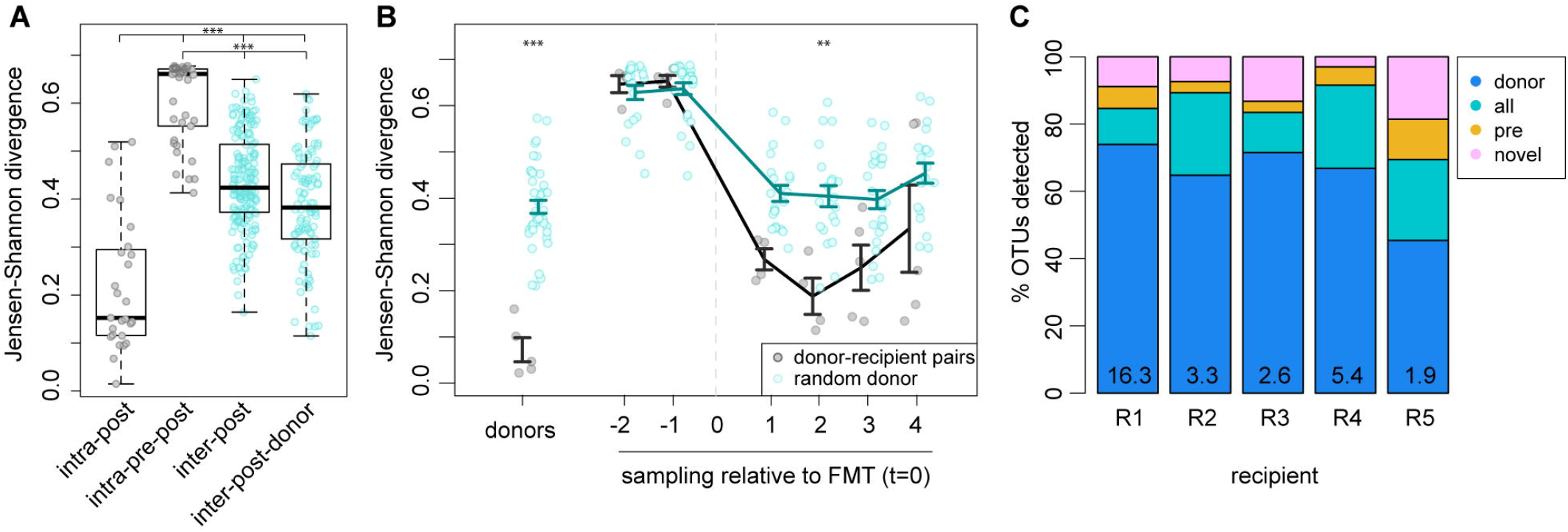
**A**) Boxplots of intra-individual community dissimilarity (Jensen-Shannon divergence) within recipients following FMT (‘intra-post’), within individuals pre- and post-FMT (‘intra-pre-post’) compared to inter-individual community similarity across patients following FMT (‘inter-post’) or compared to donors (‘inter-post-donor’). **B**) Community dissimilarity (Jensen-Shannon divergence) across time in donors and in recipients over time, compared respective donors (blue) or other donors (grey). Lines represent mean and standard deviation. (Wilcoxon test, **p < 0.005, ***p < 0.0005) **C**) Percent of total operational taxonomic units (OTUs) detected in each recipient in all samples collected post-FMT (present/absent) that were detected in their pre-FMT samples (‘pre’), respective donor (‘donor’), both pre-FMT and donor (‘all’), or detected only post-FMT (‘novel’). The percent of OTUs in each recipient that were only found in their respective donor and recipient post-FMT, and never in other individuals, is listed at the bottom of each bar.

We next calculated whether the origin of specific OTUs could be determined across donor-recipient pairs. We observed that on average, recipients shared 64.5% of total OTUs post-FMT with their respective donor that were never detected in the recipient pre-FMT (‘donor’ origin, Fig. 2C). A mean of 19.2% of OTUs in the recipients post-FMT could be detected in both donor and the recipient pre-FMT (‘all’), and a mean of 10.2% of OTUs in recipients post-FMT were present in the recipient pre-FMT and never detected in their respective donor (‘pre’). We also detected a mean of 10.2% of OTUs in recipients post-FMT that were never detected in either the recipient prior to FMT or in the donor (‘novel’). Interestingly, a mean 5.2% of total OTUs present in recipients at a given time point post-FMT that were shared with their respective donor were never detected in any other subject. Classification of these unique OTUs encompassed multiple phyla, although many unique OTUs were classified within the Clostridiales class (Table S2). These results suggest that diverse microbiota members have the capability to directly colonize from the donor, as discriminated by the presence of unique OTUs within donor-recipient pairs.

### Microbiota-derived metabolites are increased following fecal microbiota transplantation

To investigate changes in bile acids within the gut microbiota of FMT-treated individuals, we measured the concentration of primary and secondary bile acids in donor and recipient fecal samples. We observed a significant shift in the bile acid profile in recipients following FMT (PERMANOVA, p < 0.005) (Fig. S2). Following FMT, we observed a decrease in the taurine-conjugated primary bile acids taurocholic acid (TCA) and taurochenodeoxycholic acid (TCDCA), unconjugated cholic acid (CA) (Fig. 3) and some glycine-conjugated primary bile acids (Table S1). In contrast, we observed an overall increase in the secondary bile acids deoxycholic acid (DCA), lithocholic acid (LCA), and ursodeoxycholic acid (UDCA) following FMT (Fig. 4). Interestingly, increases in secondary bile acids surpassed the levels observed in donors. The concentration of secondary bile acids detected in recipient stool over the recovery period varied drastically from one recipient to another and per time point, suggesting that recovery of microbial derived secondary bile acids is a dynamic process.

**Figure 3.**
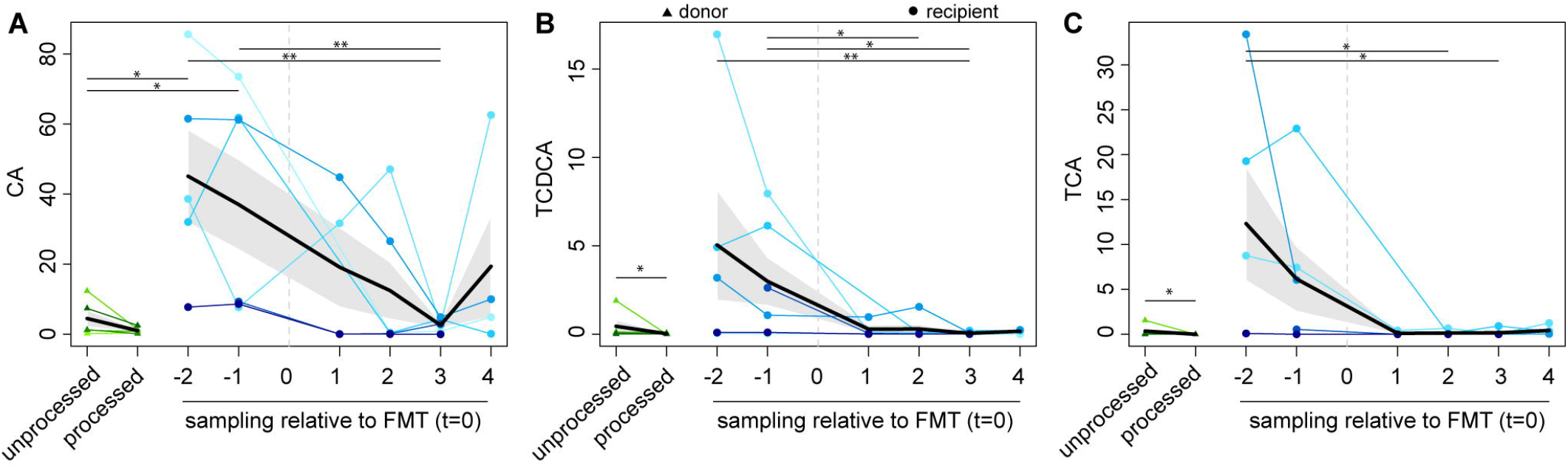
Levels of the primary bile acids **A**) cholic acid (CA), **B**) taurochenodeoxycholic acid (TCDCA), and **C**) taurocholic acid (TCA) in donors (green: unprocessed versus processed donor material) and recipients over time (blue). Individual points are shown in μg/100 mg fecal dry weight, with mean value per sampling time point (black) and standard deviation (grey area).

**Figure 4.**
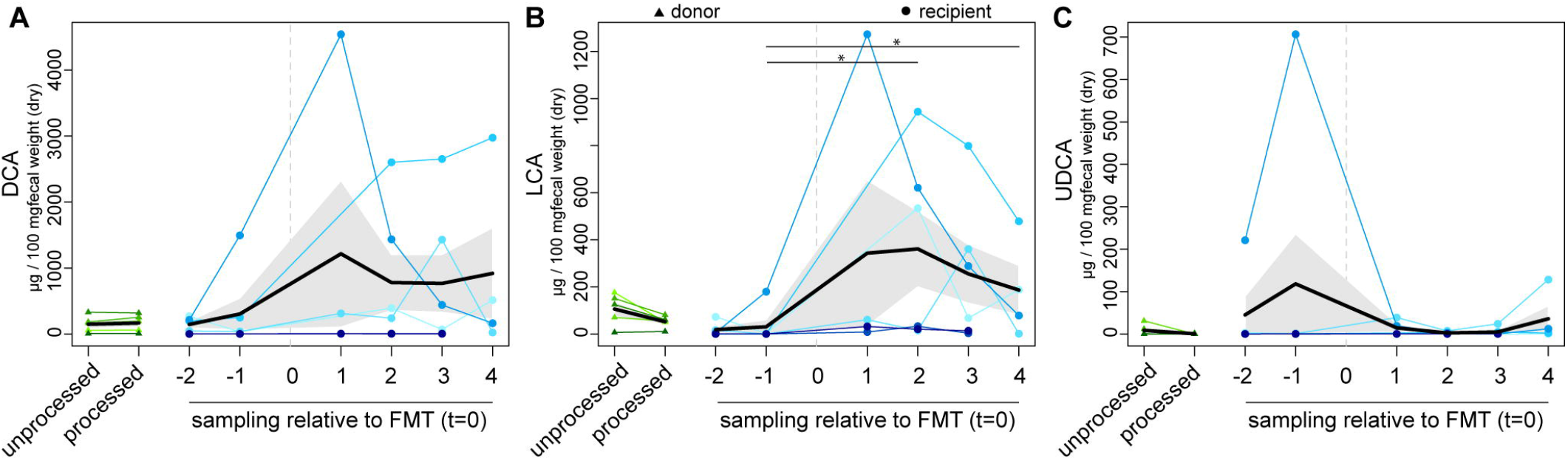
Levels of the secondary bile acids **A**) deoxycholic acid (DCA), **B**) lithocholic acid (LCA), and **C**) ursodeoxycholic acid (UDCA) in donors (green: unprocessed versus processed donor material) and recipients over time (blue). Individual points are shown in μg / 100 mg fecal dry weight, with mean value per sampling time point (black) and standard deviation (grey area).

Given the changes observed in the microbiota community, we hypothesized that the production of other microbial metabolites in addition to bile acids would increase following FMT. We observed that levels of the major SCFAs acetate, propionate, and butyrate increased in all patients treated with FMT, reaching similar levels as observed in the donor (Fig. 5). Processing the fecal material (homogenization and filtering) decreased levels of the SCFAs measured. Levels of succinate and lactate, which can be used by some bacteria as substrates for the production of propionate and butyrate respectively, remained unchanged in most patients treated with FMT (Fig. S3).

**Figure 5.**
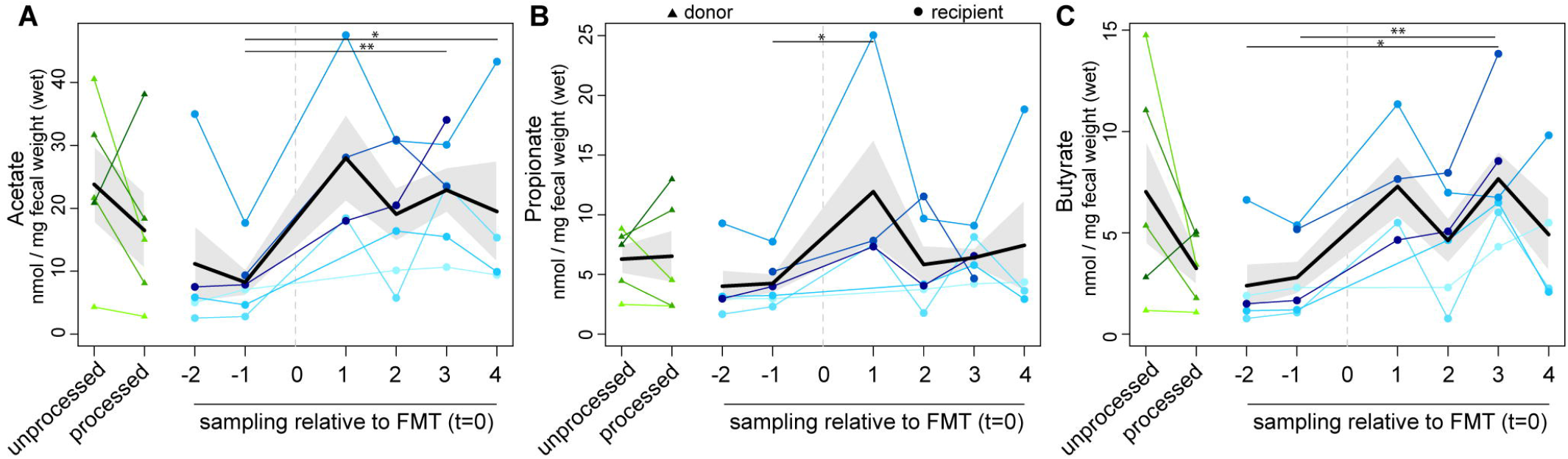
Levels of the short chain fatty acids **A**) acetate **B**) propionate, and **C**) butyrate in donors (green: unprocessed versus processed donor material) and recipients over time (blue). Individual points are shown in nmol/mg feces (wet weight), with mean value per sampling time point (black) and standard deviation (grey area).

### Microbial metabolites are correlated with diverse OTUs following fecal microbiota transplantation

Although we observed generalizable patterns of microbial community and metabolite recovery in patients receiving FMT, intra-individual similarity was higher compared to inter-individual similarity following FMT, suggesting some individuality of microbiota recovery over time (Fig 2A). To identify OTUs that associated with the levels of individual metabolites, while accounting for clustered (intra-individual), repeated measurements over time, we built multivariable mixed models using generalized estimating equations (GEEs) (25). This approach takes into account longitudinal changes within subjects while still allowing one to make global inferences about the effect of specific OTUs on metabolite concentrations across individuals. We focused on metabolites that were most impacted by FMT within recipients over time: the primary bile acids CA, TCA, TCDCA, the secondary bile acids DCA and LCA, and the SCFAs acetate, propionate, and butyrate. In total, 180 OTUs significantly associated with one or more metabolites, classified across 30 different families in six phyla (p value < 0.0001 cutoff, Table S3). OTUs correlated with metabolites encompassed broad bacterial classifications, including within the Firmicutes phylum (134 OTUs), within the Lachnospiraceae (42 OTUs), Ruminococcaceae (26 OTUs), and unclassified Clostridiales (24 OTUs) families. Most of these OTUs were positively correlated with SCFAs and secondary bile acids and negatively correlated with primary bile acids (Fig. 6). A cluster of OTUs classified within the Veillonellaceae family (Firmicutes) and several OTUs classified as Proteobacteria were negatively correlated with the secondary bile acid LCA.

**Figure 6.**
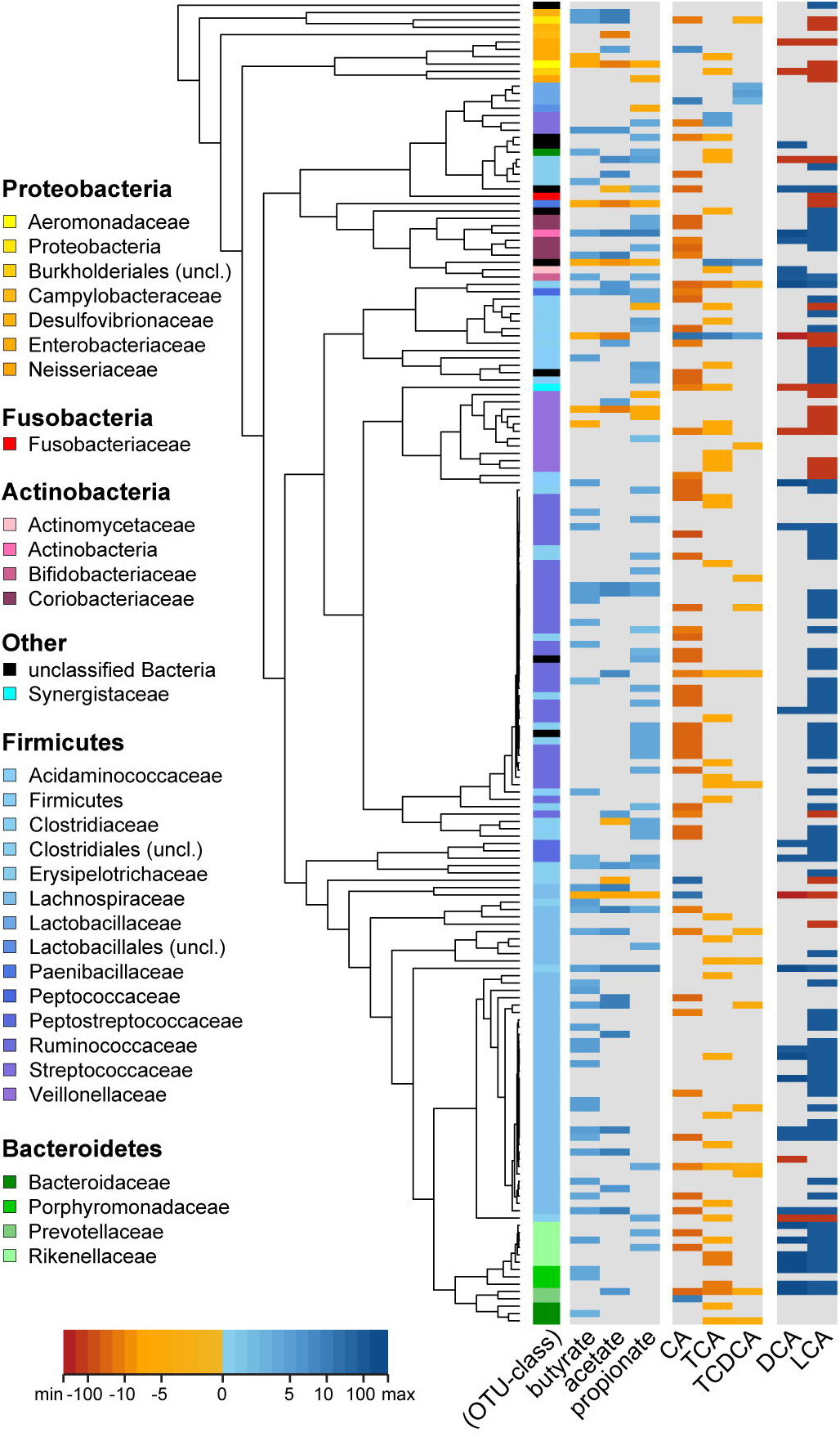
Phylogenetic tree of operational taxonomic units (OTUs) (rows) that were significantly correlated with specified short chain fatty acids and bile acids (heatmap columns; red = negative correlation and blue = positive correlation) using generalized estimating equation (GEE) models in recipients over time. The genus-level classification is indicated by the first column and accompanying legend on the right.

## Discussion

Our results demonstrate individualized recovery of the microbiome environment following FMT for the treatment of recurrent CDI. While overall microbiota diversity was increased in all patients, inter-individual differences in specific microbes were dependent on the donor and recipient, such as the detection of OTUs unique to donor-recipient pairs. In particular, we observed variable changes in bile acids over time and across patients following FMT in recipients, suggesting that secondary bile acid production throughout recovery may be more dynamic.

In our study, we observed that the majority of OTUs detected in recipients post-FMT were shared with the donor, suggesting that direct transplantation occurred. More importantly, some OTUs within individual recipients following FMT were unique to their respective donor, and never detected in other individuals, further supporting this observation. These OTUs were stably detected overtime, and encompassed a broad range of taxonomies depending on the individual. Although taxonomic resolution provided by 16S rRNA sequencing is not robust enough to distinguish strain-level differences between microbes, our results support observations made in studies that have used more discrete methods to identify bacteria that colonize following FMT. A recent study using genomes assembled from metagenomic data to distinguish more species-level colonization in two patients with CDI treated with FMT demonstrated that Bacteroidales, but not Clostridiales, were more likely to colonize individuals (26). Using single-nucleotide variation in metagenomic data to detect transfer of species following FMT in obese individuals, Li *et al*. identified that species such as *Roseburia hominis*, *Ruminococcus lactaris*, and *Akkermansia muciniphila* were able to transplant recipients from the donor (27). However, strain dominance and resilience was not consistently observed across patients, suggesting at least some individuality in acceptance of transplanted species. These studies, along with our results, support the observation that although transfer may occur during FMT, it is likely to be stochastic or donor-recipient dependent.

One of the key questions about FMT efficacy for recurrent CDI and other gastrointestinal conditions has been whether direct colonization by relevant species, or ‘engraftment’, is even necessary for successful treatment outcome. Other studies using 16S rRNA sequencing of the microbiota in patients with CDI treated with FMT have observed that colonization by donor microbes did not necessarily translate to a successful outcome (24, 28). It was also recently reported that fecal filtrate was sufficient to mediate recovery from CDI (29), which suggests a potential role for microbial metabolites in the context of the host. For non-CDI conditions, there may be more concern for matching donor-recipient microbiota communities during FMT treatment; patients with recurrent CDI exhibit a limited diversity within their microbiota compared to healthy individuals, and may be more amenable to colonization by specific microbes or a community compared to patients with IBD or obesity, who generally exhibit diverse but divergent microbiota profiles. Indeed, variable colonization has been observed in germ-free mice colonized by human microbiota (30, 31) or antibiotic-treated mice colonized by different mouse communities (32). More information not only on which microbes are relevant to each disease, but how donor-recipient dynamics impact efficacy, are likely to aid development of effective FMT therapeutics.

Regardless of the source of transplanted microbes, recovery from CDI is likely dependent on stable recovery of microbial functions that perpetuate the ability of the microbiota to provide colonization resistance against *C. difficile*. A leading hypothesis for resistance is the ability of the gut microbiota to convert primary bile acids, which include known germinants of *C. difficile* spores, into secondary bile acids, which are known to inhibit *C. difficile* growth *in vitro* (12, 16). Our data in human patients supports this trend; however, levels of secondary bile acid recovery were variable across individuals. Variation in the levels of detected bile acids following FMT in patients has also been observed in other studies both cross-sectionally and longitudinally (21, 24). Approximate levels of DCA and LCA immediately following FMT, secondary bile acids that have shown activity *in vivo* in attenuating or delaying *C. difficile* colonization in mice (20, 22), also did not predict which individuals fail FMT (24). This correlation does not necessarily discount the role of secondary bile acids in clearance of *C. difficile*, as many physiological factors, such as host response, diet, and gut transit time, can impact bile acid production at a given time point (33). It is possible that other microbial metabolites produced by the remaining community impact secondary bile acid levels, which may be more important in clearance of C. *difficile* compared to initial resistance. It is also possible that the antimicrobial properties of secondary bile acids are able to shape the recovering microbiota (34).

In addition to general shifts in bile acids, we observed full recovery of the SCFAs acetate, propionate, and butyrate following FMT. In particular, the role of butyrate is of interest given its role in maintaining the gut epithelial barrier and regulation of immunity (35, 36). Depletion of butyrate-producing bacteria has been associated with CDI in humans (37) and butyrate has been shown to limit expansion of the intestinal pathogen *Salmonella enterica* in the mouse gut (38). Little is known about the synergistic effects of bile acids and SCFAs, but the long-term recovery of SCFAs in our cohort supports persistent recovery of a metabolically active community, which may be critical to maintaining resistance to *C. difficile*. Not surprisingly, related microbes were found to correlate with increases in both bile acids and short chain fatty acids following FMT in our study. It is possible that in cases where community recovery is observed but clearance of *C. difficile* does not occur or is transient, the community is unable to function properly despite being successfully transplanted. This may be the case for *C. difficile-infected* individuals with IBD, where it has been reported that FMT has a lower success rate (39).

In summary, we observed variable direct transfer between donor-recipient pairs following successful treatment of CDI with FMT. Furthermore, recovery of the microbiome after FMT was variable across individuals both at the structural and functional levels. Resolution of dynamics of the metabolome over time, particularly of bile acids, in healthy humans could aid benchmark natural variation present in a microbiome resistant to colonization resistance against *C. difficile*, elucidating specific factors that potentially mediate clearance.

## Materials and Methods

### Patient recruitment and sample collection

Patients with recurrent *C. difficile* scheduled to receive FMT as treatment for recurrent CDI were recruited from the Infectious Diseases (ID) outpatient clinic at the University of Michigan Health System from March–December 2014. Recruited donors and recipients were over 18 and not pregnant. Recipients had experienced at least two recurrent CDI diagnoses (ascertained by documentation of symptoms by an ID physician and positive toxigenic *C. difficile* stool testing). Donors were screened by the clinical microbiology laboratory for blood-borne pathogens (4^th^ generation HIV 1/2 antibody testing, a hepatitis panel, and rapid plasma reagent for syphilis) and gastrointestinal infections (stool culture, stool ova/parasite exam, and toxigenic *C. difficile*) prior to the procedure, and all tested negative. Following informed consent, FMT recipients were asked to collect six fecal samples using a commode specimen collection hat (Fisherbrand, Cat. #02-544-208) and filling a collection tube (Sarsted Inc, Cat. #80.734.311) with fecal material: two samples prior to the procedure (1–2 weeks and 1–2 days before), and four samples following the procedure (1–2 days, 1–2 weeks, 1–2 months, and 6 months after). Samples were immediately frozen at −20°C and placed at −80°C upon retrieval of the fecal sample by a study member. Donors were asked to provide two samples at the time of the procedure: one unprocessed sample to be immediately frozen, collected as described above, and one sample processed as for the FMT procedure. For FMT procedure, donors were instructed to dilute the fecal material 1:1 with sterile saline solution, homogenize the mixture in a blender at home, and bring the sample to the hospital at room temperature. Prior to the procedure, the donor material was diluted 1:1 again with sterile saline solution. Large particular matter was then filtered out until the solution was of a consistency that could be drawn up into 50 cc syringes and administered through the accessory port of a colonoscope. Recipients underwent a standard colonoscopy bowel prep with a clear liquid diet, followed by polyethylene glycol overnight, and had nothing by mouth starting the morning of the procedure. Standard colonoscopy was performed and the FMT material was then administered via colonoscope at the level of the ileocecal valve, or as close to this area as possible. All patients had resolution of their diarrhea within 48 hours and remained symptom free until the last patient contact at 6 months post-FMT.

### DNA extractions and 16S rRNA gene-based sequencing

Samples were extracted and sequenced at the University of Michigan Microbial Systems Molecular Biology Lab, as previously described (5). Approximately 25 mg of fecal material was partitioned for DNA extraction using the MoBio PowerMag Soil DNA kit (recently renamed MagAttract PowerSoil DNA kit, Qiagen, Cat. #27100-4-EP), optimized for the epMotion 5075 TMX, using manufacturer’s directions. Briefly, PCR amplification of the 16S V4 region, using uniquely barcoded dual index primer pairs from Kozich et al (40), was conducted in a 20 μl reaction: PCR-grade water (11.85 μl), 4 μM stock combined primers (5 μl), Accuprime High-Fidelity Taq (0.15 μl) and 10X Accuprime PCR II buffer (2 μl) (Life Technologies, #12346094), with 1 μl template. PCR cycling conditions were ran at 95°C (2 min), followed by 30 cycles of (95°C (20 sec), 55°C (15 sec), and 72°C (5 min)), and 72°C (10 min). Libraries were normalized using the SequelPrep kit (Life Technologies, #A10510-01) before pooling, then quantified with the Kapa Biosystems Library Quantification kit for Illumina (KapaBiosystems, #KK4854). The Agilent Bioanalyzer high-sensitivity DNA analysis kit (#5067-4626) was used to determine amplicon size. Sequencing was conducted using the MiSeq Reagent 222 kit V2 (#MS-102-2003) (500 cycles), and 2 nM of the libraries were prepared using Illumina’s protocol, spiked with 10% PhiX for diversity for a final concentration of 4 nM. Paired-end FASTQ files for sequences used in this study are available in the Sequence Read Archive under BioProject PRJNA384621 (BioSamples SAMN06842879-SAMN06842922).

### Metabolite analysis

Targeted bile acid metabolomics was done as previously described in Theriot *et al*. (13). Briefly, for sample preparations, two-step extraction of fecal content with ethanol and 1:1 chloroform/methanol was done with isotope labeled internal standards spiked in. Extraction solvent was added to fecal samples for homogenization with a probe sonicator. Homogenized samples were kept on ice for 10 min and centrifuged at 45°C at 13000 rpm for 10 min. Supernatant containing metabolites were transferred to another tube and pellets were extracted with 1:1 of chloroform/methanol. Supernatants from the two steps were combined and dried under vacuum at 45°C by using a vacuum centrifuge. Dried samples were reconstituted in 50:50 methanol/water for LC-MS analysis. A series of calibration standards were prepared and analyzed along with samples to quantify metabolites. An Agilent 1290 liquid chromatography (Agilent, Santa Clara, CA), waters Acquity BEH C18 column with 1.7 μm particle (2.1 × 100 mm, Waters, Milford, MA) was used for chromatographic separation. Mobile phase A (MPA) was 0.1% formic acid in LC-MS grade water. Mobile phase B (MPB) was 0.1% formic acid in acetonitrile. Mobile phase gradient consisted of 0–0.5 min of 5% MPB; 3 min of 25% MPB; 17 min of 40% MPB; and 19–21 min of 95% MPB and was run on an Agilent 6490 triple-quadrupole mass spectrometer (Agilent, Santa Clara, CA). Data was processed by MassHunter workstation software, version B.06.

Targeted SCFA metabolomics was done as follows: Mouse fecal pellets were homogenized in a bullet blender (no beads), in 400 μL of a solution of 30 mM hydrochloric acid plus isotopically-labeled acetate (0.125 mM), butyrate (0.125 mM), and hexanoate (0.0125 mM.) 250 μL of Methyl tert-butyl ether (MTBE) was added to each sample, and the mixture was vortexed for 10 seconds to emulsify, then held at 4 °C for 5 mins, and vortexed again for 10 seconds. Samples were centrifuged for 1 minute to separate the solvent layers and MTBE was then removed to an autosampler vial for GC-MS analysis. 10 μl of MTBE was removed from each sample and pooled in a separate autosampler vial for quality control purposes. A series of calibration standards were prepared along with samples to quantify metabolites. GC-MS analysis was performed on an Agilent 69890N GC −5973 MS detector with the following parameters: a 1 μL sample was injected with a 1:10 split ratio on a ZB-WAXplus, 30 m × 0.25 mm × 0.25 μm (Phenomenex Cat#7HG-G013-11) GC column, with He as the carrier gas at a flow rate: 1.1 ml/min. The injector temperature was 240 °C, and the column temperature was isocratic at 310°C. Data were processed using MassHunter Quantitative analysis version B.07.00. SCFAs were normalized to the nearest isotope labeled internal standard and quantitated using 2 replicated injections of 5 standards to create a linear calibration curve with accuracy better than 80% for each standard.

### Data Analysis

Steps for data processing, analysis and figure generation are described in detail at: www.github.com/aseekatz/UMFMT. Mothur v1.37.6 (41) was used to process sequences and conduct specified downstream analyses, with UCHIME to remove chimeras (42). The SILVA rRNA database project (v119), adapted for use in mothur, was used to align sequences to the 16S V4 region (43). Operational taxonomic units (OTUs) were calculated using the average neighbor algorithm in mothur at a 97% similarity cutoff. We used the Ribosomal Database Project (RDP, v10, adapted for use in mothur) to classify OTU sequences and phylotypes using the Wang method with an 80% similarity minimum and bootstrapping (44). Alpha diversity metrics (inverse Simpson index), beta diversity metrics (Jensen-Shannon divergence), and principal component axes were calculated in mothur. Remaining analyses and visualizations were generated using R (R Foundation for Statistical Computing, Vienna, Austria, v3.1.0) and specified R packages. Standard R commands were used to summarize and visualize donor similarity measures (both intra- and inter-individual comparisons, from Jensen-Shannon divergence), diversity metrics, shared OTU percentages in donor-recipient pairs, and abundance of metabolites. For calculation of shared OTUs between recipients and their respective donors, an OTU was considered ‘present’ if detected in any of the donor, pre-FMT, or post-FMT samples within that individual at the specified time points (pre- vs. post-FMT and compared to donor). A heatmap of the relative abundance of OTUs and metabolite coefficients was generated using gplots (45). A phylogenetic tree of OTUs that were significantly correlated with select metabolites was generated by aligning the specified OTUs in mother using the SILVA database, building a phylogenetic tree of this alignment in RaXML (46), and visualized in R with metabolite coefficients using the R packages phangorn (47), phytools (48), plyr (49), and gplots.

### Statistics & modeling

Summary statistics for all measures were initially calculated using the R package dplyr (50). The nonparametric Wilcoxon (two-group comparisons) or Kruskal-Wallis test (multi-group comparisons) was used to determine significance between different treatment groups. PERMANOVA (R package vegan) was used to determine significant clustering between groups (51). In order to account for the clustered, repeated measures inherent in the dataset, generalized estimating equation (GEE) models were constructed to evaluate for associations between each OTU's absence/presence and metabolite concentrations over time. GEE models were constructed in R using the package geepack (52) and an autoregressive1 correlation structure. Models were built as follows: 1) unadjusted models of each individual OTU and the metabolite being evaluated were run to generate p values and coefficients; 2) the OTUs were ranked by most to least significant p value followed by the largest coefficient (effect size); and 3) the final multivariable GEE model for each metabolite was then built through stepwise addition of each OTU to the model, starting with those ranked highest. Criteria for retention of an OTU at each step included the quasi-likelihood ratio (QIC) calculated with the R package QICpack (53), the p values and point estimates for each OTU as the model was built, and whether the model could converge (given the small sample size, larger models utilizing more than 5–10 variables were often rank deficient). When multiple possible final models were possible, the model with the lowest QIC was used. Annotated code for analysis and generation of results are deposited in the cited github link.

## Acknowledgments

The work on this paper utilized Metabolomics Core Services supported by grant U24 DK097153 of NIH Common Funds Project to the University of Michigan. This research was supported in part by The University of Michigan Medical School Host Microbiome Initiative (HMI). We thank Judith Opp, April Cockburn, and Harriette Carrington in the HMI Microbial Systems Molecular Biology Laboratory for sample processing and sequencing. VBY has previously consulted for Merck and Vedanta Biosciences. VBY and AMS have previously received a research grant from MedImmune. CMT is a scientific advisor to Locus Biosciences, a company engaged in the development of antimicrobial technologies.

## Supplemental Figure Legends

**Table S1.** Sample info and processed sequencing results. Sample metadata (sample ID, patient ID, sampling order (time point), treatment group (donor, pre-FMT, post-FMT), sample processing (homogenized or not), relative sampling day to FMT, age, sex, donor relatedness or donor shared living) are listed in columns A-J. Basic sequencing metrics are listed in columns K-M. Samples with less than 5,000 sequences were excluded from analyses. The relative abundance of the most abundant OTUs (minimum abundance of 0.0001% in at least one sample), with classification to the genus-level, are listed in columns N-JZ. Raw normalized metabolites are listed in columns KA-LI: bile acids were measured in μg/100 mg fecal weight (dry), short chain fatty acids were measured in nmol/mg fecal weight (wet), and succinate and lactate were measured in μM/mg fecal weight (dry).

**Table S2.** Unique OTUs between donor-recipient pairs. List of OTUs and their classifications shared between recipients (post-FMT) and their respective donors that are never detected in the remaining dataset. An OTU was considered 'shared' between the pairs based on presence (versus absence) in at least one of the post-FMT samples and at least one of the donor samples sequenced.

**Table S3.** Metabolite-OTU correlations. Operational taxonomic units (OTUs) that significantly correlated with the specified metabolites (p < 0.001), with taxonomic classification to the genus level (RDP v.10).

**Figure S1.** Heatmap of the relative abundance of the top 40 most abundant operational taxonomic units (OTUs) (columns) detected in all donor and recipient samples (rows).

**Figure S2.** Nonparametric multidimensional scaling (NMDS) of Horn-Morisita dissimilarity index of detected bile acids (see Table S1) in fecal samples of recipients (pre-FMT = yellow; post-FMT = blue) and donors (green). Metabolite ordinations are labeled in red. Pre-FMT, post-FMT, and donor samples clustered significantly from each other (Analysis of Variance, or Adonis, p < 0.005).

**Figure S3.** Levels of the short chain fatty acids A) lactate and B) succinate in donors (green: unprocessed versus processed donor material) and recipients over time (blue). Individual points are shown in μM/mg fecal weight (dry), with mean value per sampling time point (black) and standard deviation (grey area). Significantly different comparisons are marked with a star (Wilcoxon test, p < 0.05).

